# Enzymatic extracellular matrix remodeling in *ex vivo* slices of aged myocardium

**DOI:** 10.64898/2026.10.19.760075

**Authors:** Ranmadusha M. Hengst, Erik M. Spaans, Md. Faris H. Ramli, Avery Rui Sun, Vicky Lai, Kim Whye Leong, Yinling Zhu, Roger S.Y. Foo, Jin Zhu, Chii Jou Chan, Leo Hwa Liang, Jennifer L. Young

**Affiliations:** Mechanobiology Institute, National University of Singapore; Singapore; Solar Energy Research Institute of Singapore, National University of Singapore; Singapore; Department of Electrical and Computer Engineering, National University of Singapore; Singapore; Department of Biomedical Engineering, National University of Singapore; Singapore; Stanford Cardiovascular Institute, Stanford University School of Medicine, Stanford, CA, USA; Department of Materials Science and Engineering, Stanford University, Stanford, CA, USA; Cardiovascular Metabolic Disease Translational Research Programme, Yong Loo Lin School of Medicine, National University of Singapore, Singapore, Singapore; Cardiovascular Research Institute, National University of Singapore; Singapore; Department of Physiology, Yong Loo Lin School of Medicine, National University of Singapore, Singapore, Singapore; Department of Biological Sciences, National University of Singapore; Singapore

**Keywords:** collagenase, stiffness, mechanics, cardiac tissue, aging, living myocardial slices

## Abstract

Cardiac aging is accompanied by extracellular matrix (ECM) remodeling and increased myocardial stiffness, which may contribute to age-associated functional decline. However, whether direct remodeling of the aged cardiac ECM can modify myocardial mechanics and function remains poorly understood. Here, we established an *ex vivo* living myocardial slice platform to investigate whether enzymatic remodeling of the aged ECM can alter myocardial structure, mechanics, and function. Left ventricular myocardial slices (LMS) from young and aged mice were maintained under static stretch for 48 hours using a custom 3D-printed culture setup. Aged slices were treated with collagenase and subsequently characterized using quantitative imaging, tensile testing, contractility measurements, and transcriptomic analysis. Aged myocardium exhibited increased collagen and fibronectin abundance, altered collagen network architecture, and increased stiffness compared with young tissue. Collagenase treatment reduced collagen and network complexity and reduced myocardial stiffness toward young tissue values without compromising ultimate tensile stress. Despite these structural and biomechanical changes, age-associated differences in myocardial contractility remained comparable across groups. Transcriptomic analysis indicated that treated aged tissue largely retained its age-associated molecular phenotype, despite pathway-level shifts involving metabolism, cardiac function, and inflammatory responses. These findings demonstrate that enzymatic ECM remodeling can modify the aged myocardial mechanical environment while selected functional and transcriptional features of aging persist over 48 hours. The LMS platform provides a controllable *ex vivo* tissue-level approach for investigating ECM contributions during cardiac aging and evaluating strategies for ECM engineering.

**Statement of significance:** Cardiac aging is accompanied by progressive extracellular matrix (ECM) remodeling and tissue stiffening, which are associated with impaired cardiac function. However, studying how these age-related changes influence living myocardium requires models that balance physiological complexity and experimental control. This study combines living myocardial slice culture with controlled enzymatic ECM remodeling to examine aged tissue responses within the native myocardial environment. Collagenase treatment reduced tensile modulus and altered collagen network organization, while selected functional and transcriptional features of aging persisted over 48 hours. These findings demonstrate the utility of living myocardial slices for evaluating matrix-targeted interventions and highlight the importance of assessing biological responses alongside changes in ECM structure and tissue mechanics.

## 1. Introduction

Aging is accompanied by progressive biological and biomechanical changes across tissues, reducing the ability of organs to adapt to physiological stress or recover from injury [1,2]. Cardiovascular disease remains a leading cause of morbidity and mortality worldwide, with the risk of heart failure increasing with age [3]. Understanding the mechanisms that drive age-related cardiac dysfunction may therefore inform strategies to preserve myocardial function.

A defining feature of cardiac aging is the progressive remodeling of the cardiac extracellular matrix (ECM), which can involve greater collagen content, altered collagen composition and crosslinking, and increased tissue stiffness [4–7]. Age-associated cardiac fibrosis can contribute to reduced ventricular compliance, impaired diastolic function, and reduced contractile efficiency [5,8–11]. These structural changes coexist with cardiomyocyte hypertrophy and reduced regenerative capacity, alongside inflammation, senescence, altered calcium handling, and changes in the sarcomeric protein titin [12–15].

The cardiac ECM, composed primarily of collagens, laminins, fibronectin, and proteoglycans, provides structural support while actively regulating cell behavior through biochemical and mechanical cues [6,11,16,17]. Alterations in ECM composition, ligand density, or spatial organization can influence cardiomyocyte and fibroblast signaling, fate determination, and mechanotransduction [17–19]. Cardiac aging has been associated with increased fibrillar collagen content and shifts in the collagen type I:III ratio in several species leading to thicker fibers [4,9,20,21]. Notably, transcript levels of major ECM components are often unchanged or reduced with age [22], highlighting the potential contributions of altered ECM turnover and post-transcriptional regulation to collagen accumulation [9,23–25]. In parallel, enzymatic crosslinking by lysyl oxidase (LOX) family enzymes [26], nonenzymatic crosslinking associated with advanced glycation end products (AGEs) [27], and the matricellular protein secreted protein acidic and rich in cysteine (SPARC) [28] can further alter myocardial mechanics.

ECM mechanics are central regulators of cardiac cell behavior. *In vitro* studies have demonstrated that cardiomyocytes and fibroblasts are mechanosensitive, exhibiting stiffness-dependent changes in morphology, contractility, and activation state. Increased substrate stiffness can promote phenotypes associated with aging and fibrosis [6,29–31], whereas matrix softening can partially reverse selected stiffness-induced responses, depending on cell type and prior exposure to stiffness [6,32–34]. These observations suggest that ECM stiffening can reinforce cellular phenotypes that contribute to cardiac dysfunction.

Beyond stiffness alone, age-associated changes in ECM composition and ligand presentation can also shape cellular phenotypes. Aged cardiac ECM has been shown to promote aging-associated phenotypes in human induced pluripotent stem cell-derived cardiomyocytes [35], while ligand presentation from young ECM can promote quiescence in cardiac fibroblasts, even when substrate stiffness mimics aged tissue [6]. Together, these findings indicate that the cardiac ECM can influence cellular aging phenotypes through interconnected biochemical, architectural, and mechanical cues. Whether modifying these cues within native aged tissue can alter myocardial function remains an important question.

Existing approaches to address cardiac fibrosis include inhibition of profibrotic signaling, modulation of matrix turnover, and interventions targeting collagen crosslinks [27,36]. Collagenase has been used experimentally and clinically to remodel dense or fibrotic tissues [37–41]. Collagenase perturbation of myocardium has also been used to investigate the contributions of collagen to myocardial structure and mechanics [42,43]. However, the structural, mechanical, and short-term biological consequences of directly remodeling an established aged matrix within living myocardial tissue remain incompletely understood.

Addressing this gap requires experimental systems that preserve native myocardial architecture, multicellularity, and functional activity while allowing direct matrix manipulation. Living myocardial slices (LMS) are thin sections of native cardiac tissue that retain physiological multicellular organization, ECM organization, and contractility under appropriate culture conditions [44–47]. LMS also respond to pro-fibrotic [48–50], inflammatory [51], and mechanical stimuli [47,48,52], making them well suited to study cell-ECM interactions in both healthy and diseased contexts.

Here, myocardial slices from young and aged mice were used to investigate whether enzymatic remodeling of collagen in aged ECM can modify tissue structure, mechanics, and function. Collagenase-mediated ECM remodeling was combined with controlled culture using a custom 3D-printed system that maintained slices under static stretch for 2 days. We assessed whether treatment shifted myocardial mechanics toward young tissue values and whether these changes were accompanied by alterations in contractile behavior and gene expression. By linking ECM remodeling to mechanical and biological responses in native tissue, this work provides an approach to assess the potential and limitations of ECM-targeted interventions in cardiac aging.

## 2. Methods

### 2.1 Tissue handling and LMS preparation

Healthy young (1-2 months) and aged (17-25 months) female C57BL/6J mice (N >= 3 each for young, aged, and treated aged, with N specified in the figure captions) were obtained through the NUS Animal Tissue Sharing Program and used in accordance with protocols approved by the Institutional Animal Care and Use Committee (IACUC) of the National University of Singapore. Young female ICR mice (1-2 months, N = 3) were additionally obtained through the NUS Animal Tissue Sharing Program for holotomography validation (Fig. 1c). Mice were euthanized by carbon dioxide asphyxiation followed by cervical dislocation.

**Fig. 1.**
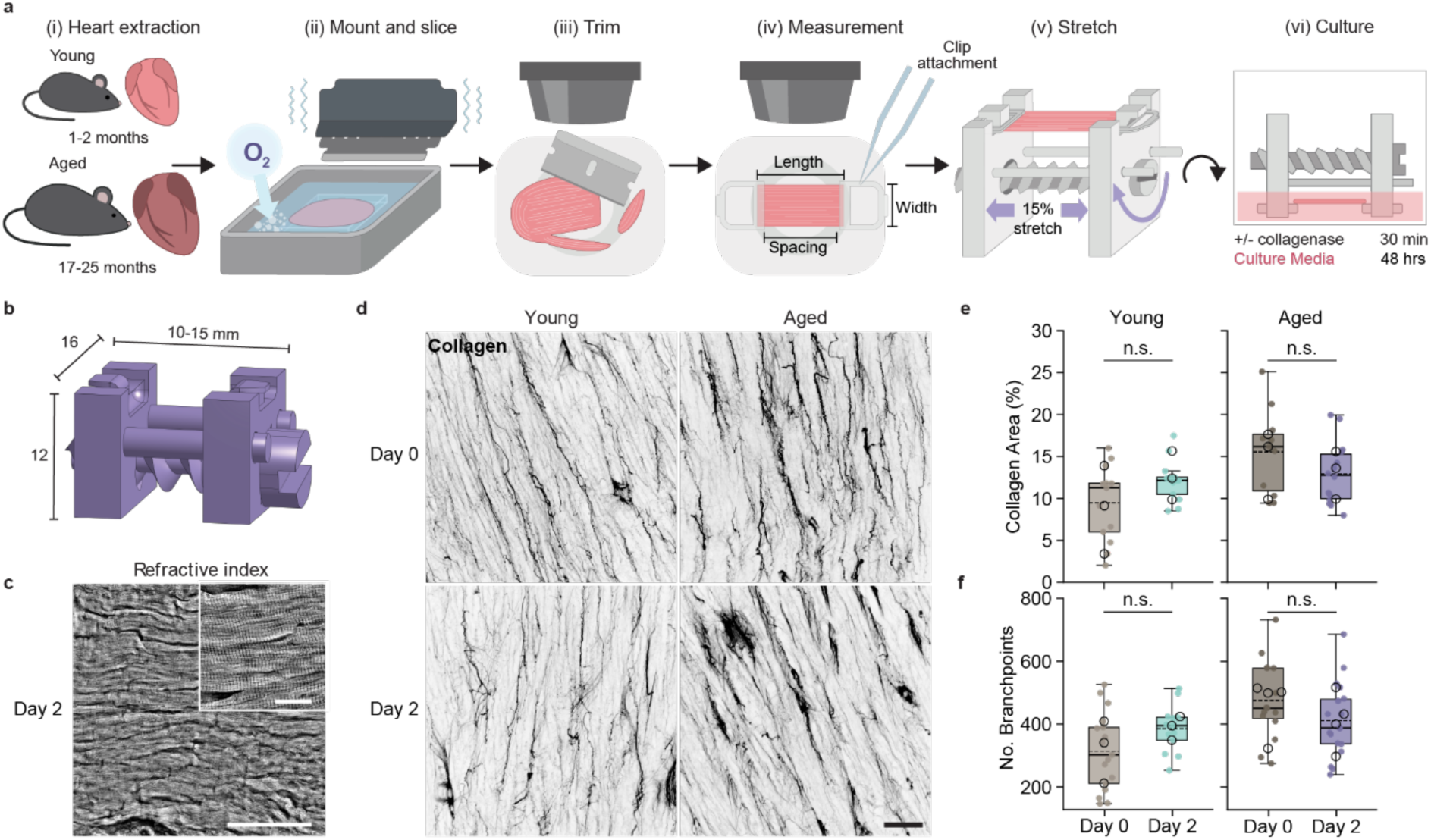
Mechanically loaded LMS culture preserves tissue organization and ECM composition over 48 hours. **(a)** Schematic overview of the living myocardial slice (LMS) isolation, mounting, and biomimetic culture workflow. **(b)** Technical schematic of the custom 3D-printed mechanical stretching device used to maintain LMS under 15% isometric stretch, with key dimensions indicated in millimeters. **(c)** Representative holotomographic images of young LMS following 48 h of culture (Day 2), showing preservation of cardiomyocyte morphology, structural alignment, and sarcomeric banding. **(d)** Representative maximum intensity confocal projections of fibrillar collagen labeled with CNA35 in young and aged LMS at baseline (Day 0) and after 48 h of culture (Day 2). **(e, f)** Quantification of collagen matrix architecture, including (**e**) collagen-positive area (%) and (**f**) number of matrix branchpoints. Box plots display the median (solid horizontal line), mean (dashed horizontal line), and interquartile range (IQR; box bounds). Whiskers extend to the most extreme data points within 1.5 × IQR of the box edges. Filled data points represent individual tissue positions (*n* = 4 − 5 fields of view per tissue); open circles indicate per-animal medians (*N* = 3 animals per group). Statistical significance was evaluated using Linear Mixed-Effects Models (LMM) with Holm-Bonferroni post-hoc correction (n.s. *p* ≥ 0.05). Scale bars = 20 *μ*m (**c,** inset) and 100 *μ*m (**c, d**).

Hearts were rapidly excised and maintained in ice-cold, sterile-filtered (0.22 µm) modified Tyrode’s solution (140 mM NaCl, 6 mM KCl, 1 mM MgCl₂, 1.8 mM CaCl₂, 10 mM HEPES, 1 mM glucose, and 30 mM 2,3-butanedione monoxime [BDM], pH 7.4) until further processing. The left ventricle was sectioned into 250-μm-thick slices using a vibratome (VT1200S, Leica Microsystems) equipped with an oxygenation system. For longitudinal sectioning, the epicardial surface of the ventricular block was affixed to a cutting disk using a thin layer of 4% low-melting-point agarose and cyanoacrylate adhesive (Vetbond Tissue Adhesive, 3M). Slices were obtained at an 18 ° cutting angle, vibration amplitude of 1 mm, and blade speed of 0.08 mm/s.

Freshly isolated slices were trimmed into rectangular specimens aligned with the predominant myocardial fiber orientation and secured at both ends to custom-designed 3D-printed clips using cyanoacrylate adhesive. The relaxed clip-to-clip distance was measured using a digital stereomicroscope (Nikon instruments), and the required extension was calculated to achieve 15% isometric stretch, maintained at fixed length during culture using a custom-designed 3D printed stretcher. Slices were mounted and stretched on the custom-designed 3D-printed stretcher described in Section 2.2, after which they were left recover in ice-cold modified Tyrode’s solution at 4 °C for 30 min before subsequent processing.

### 2.2 3D printed tissue stretcher

All 3D-printed components used in this study were designed using computer-aided design (CAD) software (Onshape, Onshape Inc.), exported as STL files, and prepared for printing using PrusaSlicer (Prusa Research). Components were fabricated from Prusament Resin Model Alabaster White (Prusa Research) using an Original Prusa SL1S SPEED resin 3D printer at a layer height of 25 μm. Printed components were washed with isopropanol (IPA), followed by curing and drying using an Original Prusa CW1S Curing and Washing Machine (Prusa Research) according to the manufacturer’s instructions. Components were subsequently rinsed in IPA and incubated overnight in Milli-Q water, followed by sequential sonication in acetone, ethanol, and Milli-Q water for 10 min each. Components were then sprayed with 70% ethanol and exposed to ultraviolet (UV) light for 2 h in a biological safety cabinet before being stored in sterile modified Tyrode’s solution until use.

The tissue stretcher comprised two tissue-attachment clips, a screw, and two complementary sliding pillars. The assembly was designed to fit within a standard 6-well plate throughout the range of extension, while maintaining the tissue approximately 0.5-2 mm above the well bottom to facilitate live imaging. The overall dimensions and assembly configuration are shown in Fig. 1B.

### 2.3 LMS treatment and culture

Following isolation and mounting, LMS were washed in sterile phosphate-buffered saline (PBS) containing 3% penicillin-streptomycin (10,000 U/mL, Gibco) and equilibrated to room temperature for 15 min. Slices were then incubated for 30 min at 37 °C and 5% CO₂ in a humidified incubator in 3 mL of either Medium 199 with Earle’s salts (Gibco) as a control or Medium 199 containing collagenase type I (Gibco, cat. no. 17100017). Collagenase was administered at a dose of 0.5 U/mm² of tissue surface area, to account for differences in tissue dimensions. Following treatment, slices were washed in sterile PBS containing 3% penicillin-streptomycin for 5 min to remove collagenase before being transferred to LMS culture medium.

LMS culture medium consisted of Medium 199 with Earle’s salts, supplemented with 3% penicillin-streptomycin, 0.1% insulin-transferrin-selenium (Gibco), 10% fetal bovine serum (Gibco), 2.15 nM triiodothyronine (Sigma-Aldrich), 100 nM dexamethasone (Sigma-Aldrich), and 20 µg/mL ascorbic acid (Sigma-Aldrich). Slices were cultured for 0 h (i.e., Day 0) or 48 h (i.e., Day 2) at 37 °C and 5% CO₂ in a humidified incubator under isometric stretch using the custom-designed 3D-printed stretcher. Culture medium was replaced twice daily.

### 2.4 Holotomography

Holotomography was performed in a separate validation cohort of young female ICR mice (1-2 months, N = 3) to assess the preservation of tissue architecture following *ex vivo* culture. This cohort was used exclusively for holotomography validation, all other experiments were performed using C57BL/6J mice. Following the culture period, LMS were fixed for 20 min at room temperature in 3.7% formaldehyde (Sigma-Aldrich) while maintained in the stretched state. Holotomography images were acquired at room temperature using Tomocube HT-X1 (Tomocube Inc.) equipped with a 40x air objective (NA 0.95). Three-dimensional refractive index reconstructions were generated using the HTX processing server (Tomocube Inc.).

### 2.5 Immunofluorescence staining and confocal imaging

Fixed LMS were simultaneously permeabilized and blocked for 2 h at room temperature in 2% bovine serum albumin (BSA, Sigma-Aldrich) prepared in 0.2% Triton X-100 (Sigma-Aldrich). Slices were then incubated overnight at 4°C with primary antibodies and fluorescent probes diluted in PBS containing 2% BSA. Collagen was visualized using EGFP-CNA35, produced using the pET28a-EGFP-CNA35 plasmid (Addgene) by the Protein Production Platform at Nanyang Technological University (NTU). The probe was used at a final concentration of 2 µM from a 127 µM stock solution. Following PBS washes, slices were incubated with fluorophore-conjugated secondary antibodies for 2 h at room temperature in PBS containing 2% BSA. Nuclei were counterstained with Hoechst 33342 (Invitrogen) for 10 min. Immunostained slices were mounted on microscope slides using Fluoromount (Invitrogen) and allowed to cure for approximately 2 days before imaging. Reagent details, including suppliers, catalogue numbers, and dilutions, are provided in Supplementary Table 1.

Confocal images were acquired using a Nikon A1R laser-scanning confocal microscope (Nikon Instruments, Japan) equipped with either a 20× air objective for low-magnification imaging or a 60× water-immersion objective for high-resolution imaging. Image acquisition was performed using NIS-Elements software (Nikon Instruments). To minimize artifacts associated with tissue handling and edge effects, image acquisition was restricted to regions away from the outer tissue margins.

### 2.6 ECM image analysis

Raw confocal image stacks were imported into Fiji/ImageJ for standardized preprocessing and quantitative analysis. Background correction and intensity adjustment, where applied, were performed using identical processing parameters across experimental groups. ECM abundance was quantified from collagen and fibronectin fluorescence images using an identical analysis workflow. ECM-positive signal was segmented from background using the automatic Triangle thresholding method, and the resulting positive area was quantified using the Analyze Particles function. ECM-positive area was normalized to the corresponding actin-positive tissue area to account for differences in tissue coverage between images. Multiple non-overlapping regions of interest were analyzed for each tissue slice, and measurements were averaged at the animal level prior to statistical analysis.

ECM architecture was quantified using TWOMBLI (The Workflow Of Matrix BioLogy Informatics), an open-source Fiji/ImageJ macro for automated analysis of fibrillar extracellular matrices [53]. Maximum-projected collagen and fibronectin images were used as inputs for TWOMBLI. Analysis parameters were optimized using representative images and subsequently held constant across all experimental groups. Multiple non-overlapping regions of interest were analyzed for each tissue slice. TWOMBLI-derived parameters describing fiber properties, network connectivity, and global matrix organization were used to characterize ECM architecture. All TWOMBLI-derived parameters were used for principal component analysis (PCA).

### 2.7 Tensile mechanical testing

Uniaxial tensile testing was performed using a Biaxial BioTester (CellScale Biomaterials Testing, Waterloo, Canada). LMS were removed from the culture stretchers with the 3D-printed tissue-attachment clips intact and mounted on the tester along the predominant myocardial fiber direction. The tissue was secured between custom-designed 3D-printed grips using small strips of fine sandpaper to improve force distribution. The LMS and attached clips were immersed in room-temperature Tyrode’s solution containing BDM throughout testing. Samples were positioned from a fully slack state, and the load cell was zeroed before each measurement to establish a consistent reference state. No preconditioning was applied. Tissues were extended at a constant strain rate of 1%/s until complete failure, with force recorded at 30 Hz using a 0.5 N load cell.

### 2.8 Mechanical analysis

Force-displacement data were processed using a custom Python pipeline to generate engineering stress-strain curves. LMS dimensions were measured during tissue preparation and used to calculate the initial cross-sectional area for mechanical testing, as well as to determine the applied stretch and collagenase dosage. Specimen thickness was defined by the vibratome cutting setting (250 µm), whereas tissue width and length were measured digitally from stereomicroscope images acquired with a scale reference.. Engineering stress (*σ*) was calculated by dividing the smoothed force by the initial cross-sectional area:

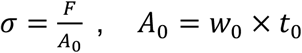

where (*F*) is the smoothed force, (*A*_0_) is the initial cross-sectional area, and *w*_0_ and *t*_0_ are the initial specimen width and thickness, respectively. Force data were smoothed using a moving-average filter with a window of 100 data points. Engineering strain (*ε*) was calculated relative to the initial gauge length:

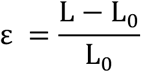

where L is the instantaneous specimen length and (*L*_0_) is the initial gauge length. Despite mechanical zeroing of the load cell prior to testing, residual slack in the mounted tissue was accounted for computationally. L_0_ was defined as the specimen length at the first point at which the smoothed force exceeded 0.2 mN, taken to indicate the onset of measurable tension following removal of residual slack. Data preceding this point were excluded from subsequent mechanical analysis.

Mechanical properties were determined from the resulting stress-strain curves using the publicly available Dots-on-Plots analysis tool [54]. The linear elastic modulus, ultimate tensile strength (UTS), strain at failure, transition point, yield point, and toughness were quantified. The transition and yield points were identified using the Dots-on-Plots algorithm with stress-deviation thresholds of 3% and 1%, respectively. The rupture point was defined as the point at which stress decreased to 70% of the ultimate stress. Data were excluded when tissue-related defects rendered the mechanical measurements unreliable, including specimens with visible damage incurred during handling or clip placement, or where tissue morphology (e.g., prominent vessels) prevented reliable mechanical characterization.

### 2.9 Contractility imaging

Contractile activity of LMS was assessed by widefield time-lapse microscopy whilst slices remained mounted on the tissue stretcher. Slices were maintained at 37 °C in Hanks’ Balanced Salt Solution (HBSS, Gibco) throughout imaging. Electrical field stimulation was applied at 1 Hz using a custom-built in-house pacing system with a peak-to-peak voltage of 20 V, a duty cycle of 1%, and a pulse width of 10 ms. Images were acquired using an Olympus Live inverted microscope with a 10× objective and controlled via MetaMorph software (Molecular Devices) at frame intervals of 10.26-50 ms, yielding recordings of approximately 500-1000 consecutive frames per field of view. Pixel binning was applied during acquisition to improve signal-to-noise ratio. For each LMS, three to four spatially distributed fields of view were acquired across the tissue surface.

### 2.10 Contractility analysis

Each image sequence was subdivided into a 4×4 grid of 16 equally sized regions of interest (ROIs), which were analyzed independently using the MUSCLEMOTION plugin for Fiji [55]. Temporal and magnitude parameters were automatically calculated per detected beat. Contractile frequency was derived as the reciprocal of the peak-to-peak interval. Normalized contraction amplitude was computed as the ratio of contraction amplitude to baseline displacement, representing a local fractional shortening analogue [56].

Quality control and data validity screening were performed in two stages prior to downstream analysis to ensure that only ROIs yielding reliable contractile measurements were included. First, tracking fidelity was evaluated by comparing the numerical derivative of the displacement trace with the MUSCLEMOTION speed trace. An ROI was retained only when all five criteria were simultaneously satisfied: (1) Pearson correlation coefficient (r) >0.7, (2) coefficient of determination (R²) >0.4, (3) regression slope >0.5, (4) root mean square error (RMSE) <0.13, and (5) mean absolute deviation (MAD) <0.1. ROIs failing all criterions were excluded on the grounds that the displacement trace could not be considered a reliable representation of contractile motion, regardless of whether the underlying biology was normal or abnormal. Second, remaining traces were screened via blinded visual inspection to exclude imaging artefacts.

Accepted parameter values were aggregated through a hierarchical structure. For each ROI, the median value across all detected beats was computed to yield a single ROI-level estimate, robust to occasional outlier beats. ROI-level values were subsequently aggregated by median across all ROIs within a given field of view to yield position-level estimates. Position-level values were further aggregated by median to yield a single animal-level estimate per condition.

Contraction traces exported from MUSCLEMOTION were additionally re-analysed using custom Python code. Traces were smoothed and baseline-corrected, and beats were identified from peaks exceeding 40 % of the typical beat size of the recording. Peaks less than 300 ms apart or separated by a valley above 50 % of the lower peak, were merged into a single beat. Each beat was then classified relative to the rhythm of its own recording: beats containing two distinct peaks (the smaller reaching at least 50 % of the larger) were classified as double peak, beats preceded by an interval shorter than 75 % or longer than 150 % of the median interval of the recording as short or long interval, respectively, and all remaining beats as regular. Double-peak, short-interval and long-interval beats were termed irregular, and an ROI was considered irregular if at least 20 % of its beats were irregular.

### 2.11 RNA extraction and sequencing

Following 48 h of culture, LMS were washed with sterile PBS to remove residual culture medium before being transferred to 1 mL TRIzol reagent (Ambion), snap-frozen in liquid nitrogen, and stored at −80 °C until RNA extraction. Frozen samples were thawed on ice and mechanically homogenized using a TissueLyser II (Qiagen). Total RNA was isolated using a TRIzol-based extraction protocol. Briefly, homogenized samples underwent chloroform-mediated phase separation with linear acrylamide as a carrier, followed by isopropanol precipitation, ethanol washing, and resuspension in molecular-grade water (Cytiva). RNA quality and integrity were assessed using an Agilent TapeStation system prior to sequencing.

Poly(A) mRNA enrichment, library preparation, and bulk RNA sequencing were performed by BGI Genomics (Hong Kong). Briefly, mRNA was enriched from total RNA using oligo(dT)-attached magnetic beads and fragmented. First-strand cDNA was synthesized using random N6 primers, followed by second-strand synthesis. The double-stranded cDNA underwent end-repair, 5’-phosphorylation, A-tailing, and adapter ligation. Following PCR amplification, the products were denatured and circularized to generate single-stranded DNA circular libraries. Sequencing was performed on the DNBSEQ platform to generate 150-bp paired-end (PE150) reads.

### 2.12 RNA-seq analysis

Primary bioinformatics processing was executed via the BGI Dr. Tom platform pipeline. Raw sequencing reads were quality-filtered using SOAPnuke (v1.5.6) to remove adapter contamination, sequences with high unknown base content (N > 0.1%), and low-quality reads. The resulting clean reads were aligned to the Mus musculus reference genome (Ensembl release 105) using HISAT2 (v2.2.1). Alignment to reference transcripts and gene expression quantification were performed utilizing Bowtie2 (v2.5.0) and RSEM (v1.3.1) to generate raw gene read counts.

Downstream transcriptomic analyses were conducted in the R statistical environment (v4.5.3). Raw read counts were imported and pre-filtered to retain genes with a cumulative count of ≥ 10 across all samples. Differential gene expression (DEG) analysis was performed using the DESeq2 package [57]. To improve the stability of effect-size estimates for genes with low counts or high dispersion, adaptive log₂ fold change (LFC) shrinkage was applied using the ashr estimator. Significantly differentially expressed genes (DEGs) were defined strictly by a Benjamini-Hochberg adjusted P-value (padj) < 0.05 and an absolute log₂ fold change > 1.

For exploratory data analysis, count data were normalized using a variance-stabilizing transformation (VST). Heatmaps of significant DEGs were generated in TBtools (v2.390; [58]). VST values were standardized by row (z-score), and genes were hierarchically clustered using Euclidean distance and complete linkage.expression values standardized across each gene using row-wise Z-scores. Functional annotation and pathway enrichment were performed using the clusterProfiler package [59], with mouse gene annotations supplied by the org.Mm.eg.db database (v3.20.0). Gene Set Enrichment Analysis (GSEA) was performed on the entire detected transcriptome to capture pathway-level changes. For GSEA, genes were strictly ranked based on the DESeq2-derived Wald statistic. Pathways were considered significant at padj < 0.05. Pathways meeting padj < 0.05 and |NES| > 1.5 were selected for visualization and assigned to functional categories, and the corresponding heatmaps were generated in TBtools.

### 2.13 Statistical tests and data visualization

All statistical analyses were performed in Python (v 3.10) using *statsmodels* (v 0.15.0) and *SciPy* (v 1.15). To account for the hierarchical structure of the data (multiple measurements acquired per animal) and avoid pseudoreplication, linear mixed-effects models (LMM) were fitted. Experimental conditions (i.e.,., Young, Aged, Treated Aged) were modeled as fixed effects, while individual animals were modeled as random intercepts. Hypothesis testing for group differences was conducted using Wald z-tests. Selected right-skewed parameters (such as ratios, coefficients of variation, and variability measures) were log-transformed for statistical testing while displayed as raw values in figures. Pairwise group contrasts were derived and *p*-values were adjusted across comparisons per variable using the Holm-Bonferroni step-down method to control family-wise error rates. Where model convergence failed or the LMM variance matrix was singular, a fallback chain sequentially utilized ordinary least squares with animal-clustered standard errors or Welch’s *t*-test on per-animal medians. Significance levels are denoted as follows: *** *p* < 0.001, ** *p* < 0.01, * *p* < 0.05, n.s. *p* ≥ 0.05. Although each aged animal contributed one control and one collagenase-treated slice, aged control and aged treated conditions were treated as independent groups for the purposes of statistical analysis.

Continuous data distributions across experimental groups are presented using box-and-whisker plots or kernel density estimation (KDE) distributions, with probability density reported on the y-axis in arbitrary units (a.u.). Box-and-whisker plots display the interquartile range (IQR), median (solid line), and mean (dashed line) of all pooled positional measurements, with individual measurements overlaid as small, filled scatter points. Per-animal medians are additionally overlaid as distinct open circles to visually differentiate technical from biological variation. For temporal and contractile profiles, representative time-series traces are displayed alongside derived stability metrics. Multi-parameter feature profiles across conditions are presented using radar and dot-plots plots, with values normalized relative to the young baseline control group. Data visualization was generated using *Matplotlib* (v 3.10) and refined in Adobe Illustrator.

## 3. Results

### 3.1 Short-term myocardial slice culture preserves tissue and ECM organization

Left ventricular living myocardial slices (LMS, 250 µm thickness) were generated from young (1-2 months) and aged (17-25 months) mouse hearts (Fig. 1a). Slices were mounted on a custom-designed 3D-printed stretcher and maintained at a 15% extension relative to the relaxed clip-to-clip distance (Fig. 1b). Static mechanical loading has previously been shown to support myocardial organization during slice culture [45]. The stretcher accommodated a range of murine LMS dimensions while remaining compatible with standard six-well culture plates and microscopy applications. Slices were maintained under these conditions for 48 hours.

To assess whether short-term culture altered cellular or ECM organization, holotomography was used to evaluate cellular morphology and sarcomere-associated banding, while immunohistochemistry was used to assess collagen and fibronectin distribution and organization (Fig. 1c-e). Following 48 hours of culture, cardiomyocytes exhibited elongated morphology and aligned organization, with characteristic repetitive banding evident by holotomography (Fig. 1c, Fig. S1c).

Fibrillar collagen was visualized using CNA35, a fluorescent collagen-binding probe that labels fibril-forming collagens, including types I and III [60]. Confocal imaging and quantitative analysis revealed no significant changes in collagen networks in either young or aged slices over the 48-h culture period (Fig. 1d,e). Specifically, collagen content (% area) and number of branchpoints revealed no significant changes over culture time (Fig. 1e). Fibronectin similarly remained unchanged in young LMS during culture (Fig. S1a,b). Thus, short-term culture did not induce detectable remodeling of the major ECM components examined. Collectively, these findings support the use of this LMS culture system for subsequent investigation of age-dependent ECM remodeling.

### 3.2 Collagenase reduces collagen and alters matrix architecture in aged slices

The effects of collagenase treatment on tissue structure and ECM organization were next assessed. LMS from aged mice were treated with collagenase for 30 min prior to culture and maintained under static stretch for 48 hours before analysis. Cells retained their elongated, aligned morphology following collagenase treatment (Fig. 2a). To assess fibrillar collagen distribution, slices were stained with CNA35 (Fig. 2b). Quantification showed that aged LMS exhibited significantly greater collagen-positive area than young LMS (Fig. 2d). Collagenase treatment significantly reduced the collagen-positive area relative to untreated aged LMS, approaching levels observed in young tissue (Fig. 2d; Fig. S2c).

**Fig. 2.**
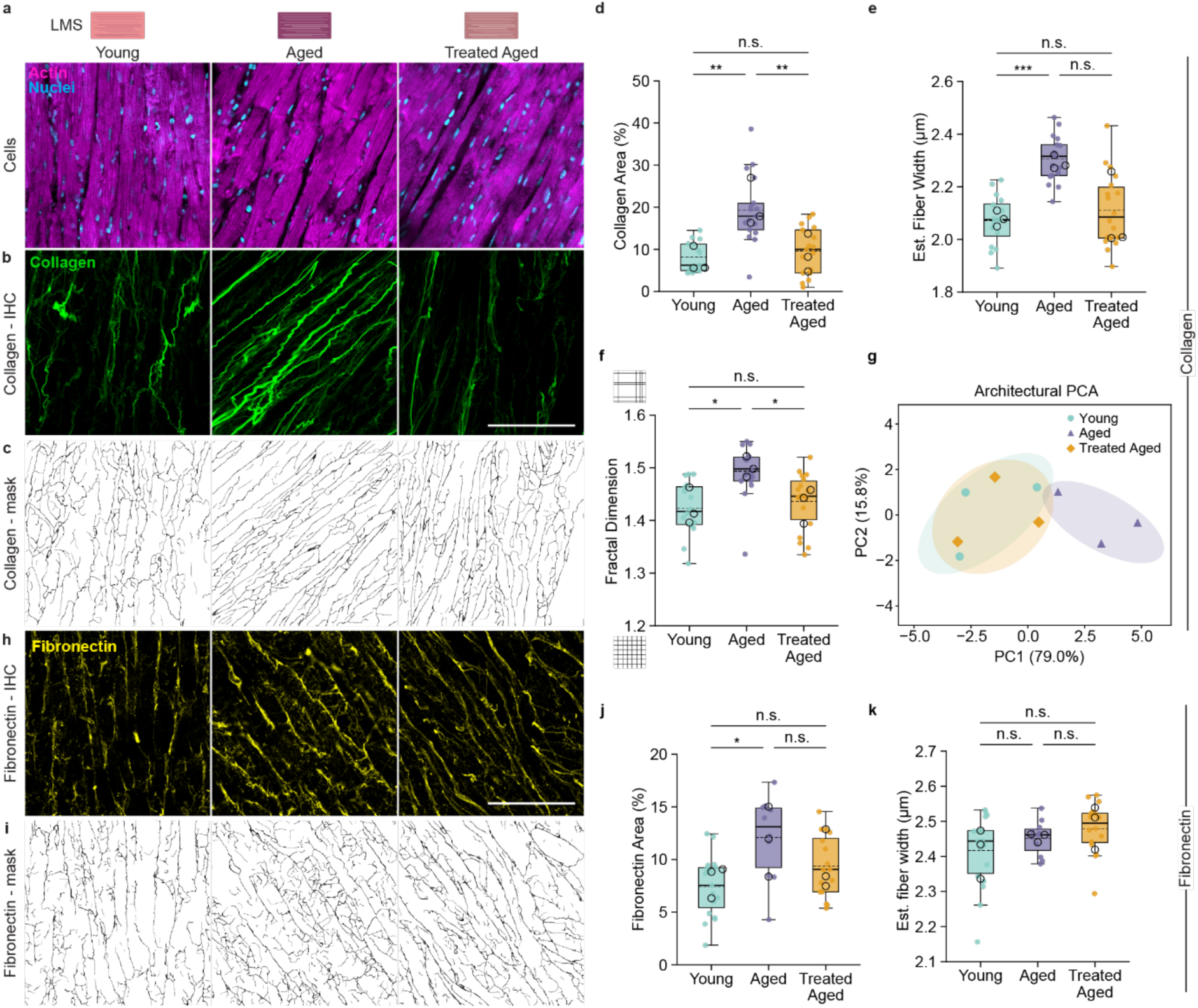
ECM remodeling following collagenase treatment of aged LMS. **(a)** Representative fluorescence confocal images of young, aged, and collagenase-treated aged LMS showing cellular organization (phalloidin/actin, magenta; Hoechst/nuclei, blue). **(b)** Representative maximum intensity confocal projections of fibrillar collagen and **(c)** corresponding TWOMBLI-generated binary matrix masks. **(d– f)** Quantification of collagen matrix architecture, including (**d**) collagen-positive area (%), (**e**) estimated collagen fiber width, and (**f**) fractal dimension (box-counting). **(g)** Principal component analysis (PCA) based on TWOMBLI-derived architectural metrics. **(h)** Representative maximum intensity confocal projections of fibronectin with **(i)** corresponding TWOMBLI matrix masks. **(j, k)** Quantification of fibronectin (**j**) positive area (%) and (**k**) estimated fiber width. Box plots display the median (solid horizontal line), mean (dashed horizontal line), and interquartile range (IQR; box bounds). Whiskers extend to the most extreme data points within 1.5 × IQR of the box edges. Filled points represent individual tissue positions (*n* = 5 − 7 fields of view per tissue); open circles indicate per-animal medians (N = 3 animals per group). Statistical significance was assessed using Linear Mixed-Effects Models (LMM) with Holm-Bonferroni correction (∗∗∗ p < 0.001, ∗∗ p < 0.01, ∗ p < 0.05, n.s. p ≥ 0.05). Scale bar = 100 µm.

To further characterize collagen organization, architectural parameters were extracted using TWOMBLI (Wershof et al., 2021) (Fig. 2c). Estimated collagen fiber width was significantly greater in aged than in young LMS, but did not differ significantly between treated and untreated aged LMS (Fig. 2e). Aged LMS also exhibited increased box-counting fractal dimension, indicative of greater network complexity, which was significantly reduced following collagenase treatment (Fig. 2f). Normalized endpoint number did not differ significantly among groups (Fig. S2b). PCA of the combined architectural descriptors revealed separation between young and aged LMS, while treated aged LMS clustered closer to young tissue (Fig. 2g). The broader set of collagen architectural descriptors is summarized relative to young LMS as a radar plot in Figure S2a.

To assess potential effects on non-collagenous ECM components, fibronectin abundance and organization were subsequently examined. Fibronectin networks were distributed throughout the slices in all three groups (Fig. 2h,i). Aged LMS exhibited a significantly higher fibronectin-positive area fraction than young LMS, consistent with reported age-related alterations in fibronectin deposition [22,35] (Fig. 2j). Following collagenase treatment, fibronectin content did not differ from that in untreated aged LMS (Fig. 2j). Similarly, estimated fibronectin fiber width and fractal dimension did not differ significantly among groups (Fig. 2k; Fig. S2d). Taken together, these data demonstrate that collagenase treatment selectively reduces collagen content and network complexity in aged LMS, shifting the collagen architectural profile toward that of young tissue.

### 3.3 Collagenase reduces myocardial stiffness in aged slices

To determine whether collagenase-mediated remodeling of the aged ECM altered myocardial mechanical properties, young, aged, and collagenase-treated aged LMS were subjected to uniaxial tensile testing (Fig. 3a). Mechanical loading was applied along the principal fiber direction to assess tissue-level passive mechanical behavior and failure properties. Stress-strain responses revealed pronounced age-dependent differences at strains of up to 25% (Fig. 3b). Aged LMS exhibited a steeper stress-strain response than young LMS, consistent with increased myocardial stiffness. Following collagenase treatment, the stress-strain response of aged LMS shifted toward that of young tissue.

**Figure 3.**
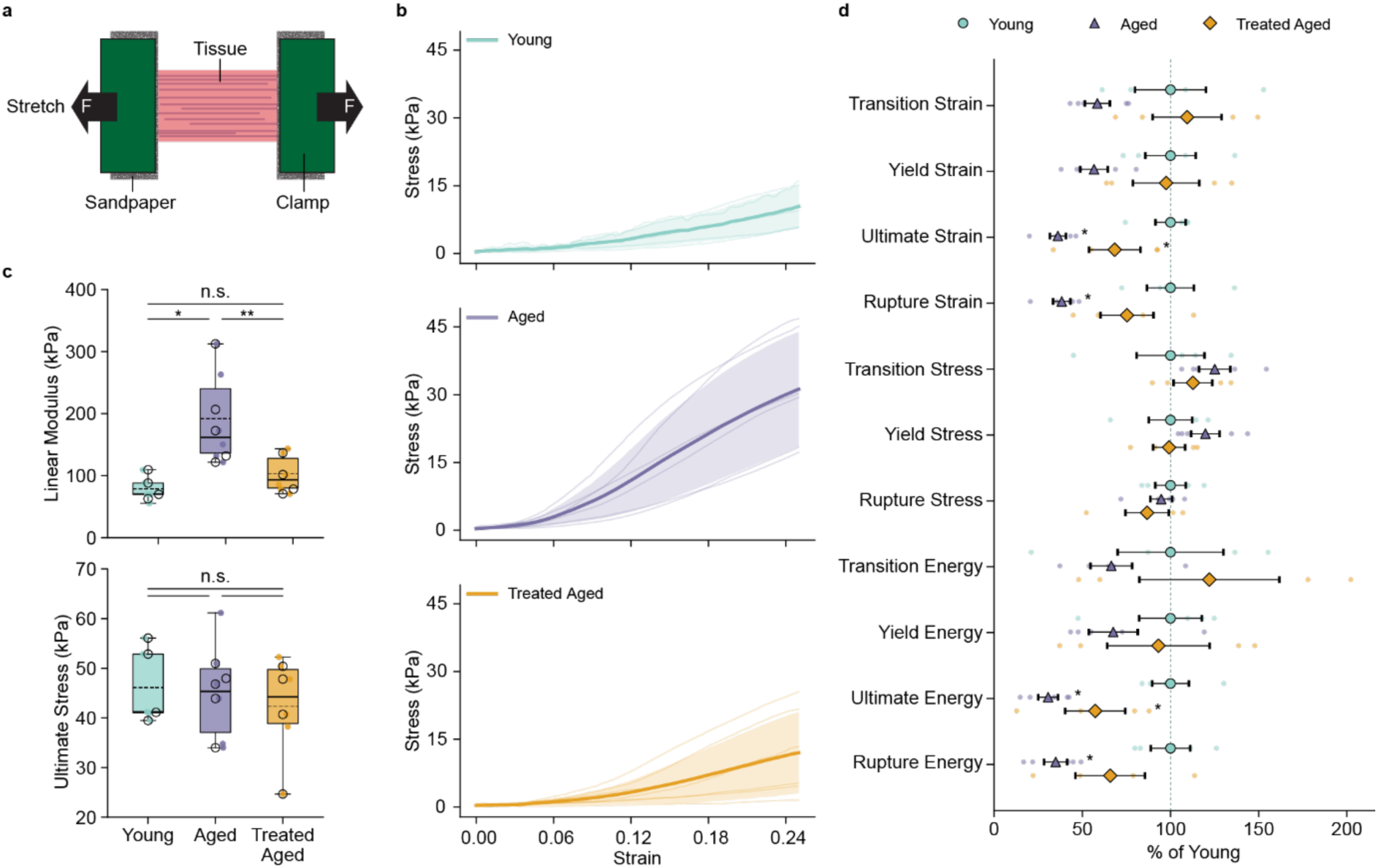
Collagenase treatment reduces the elevated stiffness of aged LMS. **(a)** Schematic representation of the uniaxial tensile testing setup. **(b)** Mean (±SD) stress-strain response curves for young, aged, and collagenase-treated aged LMS across the low-strain regime (0–25% strain); individual raw tissue curves are shown as semi-transparent overlays. **(c)** Quantification of primary passive mechanical metrics, including linear elastic modulus and ultimate tensile stress. **(d)** Relative changes in secondary tensile parameters presented as percentage change relative to young controls (mean ±SEM) with annotated significance (∗ *p* < 0.05 vs. Young). Box plots display the median (solid horizontal line), mean (dashed horizontal line), and interquartile range (IQR; box bounds). Whiskers extend to the most extreme data points within 1.5 × IQR of the box edges. Filled points represent individual LMS (*n* = 1 − 3 tissue slices per animal); open circles indicate per-animal medians (*N* = 4 young, *N* = 5 aged, *N* = 4 collagenase-treated aged animals). Statistical significance was evaluated using Linear Mixed-Effects Models (LMM) with Holm-Bonferroni correction (∗∗ p < 0.01, ∗ p < 0.05, n.s. p ≥ 0.05).

Key mechanical features, including the transition point, yield point, and linear elastic modulus, were extracted from the stress-strain curves using the Dots-on-Plots platform [54] (Fig. S3a,b), with the complete set of material parameters summarized in Supplementary Table 2. The linear modulus of aged LMS was ∼2.4 times that of young slices (192.0 ± 77.8 versus 78.6 ± 20.8 kPa, mean ± SD; Fig. 3c), confirming the increased stiffness of aged myocardium. Collagenase treatment significantly reduced the linear modulus of aged LMS by ∼47%, to 102.7 ± 30.8 kPa. The linear modulus of treated aged LMS did not differ significantly from that of young LMS.

In contrast, no significant differences in ultimate tensile stress were detected among groups (Fig. 3c). Comparison of the broader set of mechanical parameters, expressed relative to young LMS, revealed lower strain and energy at both ultimate stress and rupture in aged LMS (Fig. 3d; Fig. S3c). Following collagenase treatment, strain and energy at ultimate stress remained lower than in young LMS. Together, these results demonstrate that collagenase-mediated remodeling of the aged ECM reduces myocardial stiffness toward young tissue levels while preserving tissue strength.

### 3.4 Age-associated differences in contractility persist after collagenase treatment

To determine whether collagenase-mediated ECM remodeling altered cardiac contractile function, LMS were subjected to 1 Hz electrical pacing using an in-house pacer, and tissue motion was quantified using MUSCLEMOTION [55]. Young LMS generally followed the imposed pacing frequency, exhibiting regular contraction peaks with consistent amplitude and duration (Fig. 4a). In contrast, aged LMS displayed greater heterogeneity in contractile behavior, characterized by irregular peak morphology, variable contraction amplitudes, and additional contraction peaks. Collagenase-treated aged LMS exhibited both regular and irregular contraction patterns.

**Figure 4.**
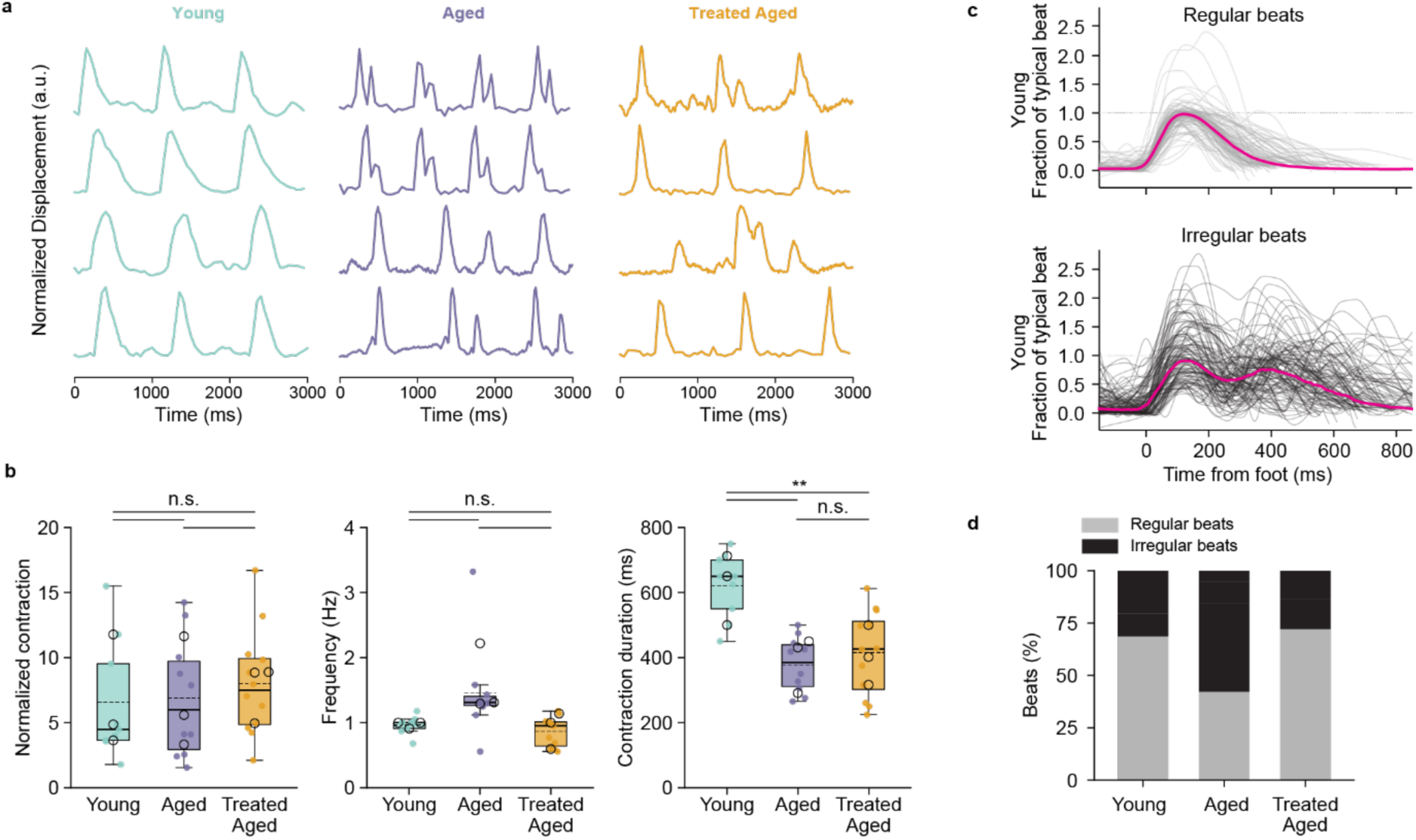
Contractile behavior of young, aged, and collagenase-treated aged LMS. **(a)** Representative contraction traces during 1 Hz field stimulation (four traces per group, 3000 ms window, each normalized to its own range). Traces were selected as the recordings closest to the group median. **(b)** Normalized contraction (ratio of contraction amplitude to baseline displacement), contractile frequency (reciprocal of the peak-to-peak interval) and contraction duration (as defined by MUSCLEMOTION). Box plots display the median (solid horizontal line), mean (dashed horizontal line), and interquartile range (IQR; box bounds). Whiskers extend to the most extreme data points within 1.5 × IQR of the box edges.. Filled points represent individual positions (*n* = 3 − 4 fields of view per slice, calculated as medians across 16 ROIs per field) and open circles per-animal medians (*N* = 3 animals per group). Groups were compared pairwise using linear mixed-effects models with animal as random intercept (Wald z), and p-values were adjusted per parameter across the three comparisons by the Holm–Bonferroni method (∗∗ *p* < 0.01, n.s. *p* ≥ 0.05). **(c)** Aligned and scaled beats (fraction of the typical regular beat of each recording) classified automatically as regular (top) or irregular (bottom) for young LMS. **(d)** Percentage of regular and irregular beats per group, calculated per animal and averaged across animals (*N* = 3 animals per group; total beats: young = 3301, aged = 1920, treated aged = 2168). Differences between groups were not significant (Welch’s t-test on per-animal percentages, Holm-adjusted (*p* ≥ 0.51).

Analysis of the three selected contractile parameters showed no significant differences in normalized contraction amplitude or frequency among groups (Fig. 4b). In contrast, contraction duration was significantly greater in young LMS than in both untreated and collagenase-treated aged LMS, with no significant difference between the two aged groups.

Beats were further classified as regular or irregular (Fig. 4c). The proportion of regular beats was lower in aged LMS and higher following collagenase treatment, but differences among groups were not statistically significant (Fig. 4d). Taken together, these findings indicate that remodeling of the aged ECM with collagenase was accompanied by a high proportion of regular beats, while age-associated differences in contraction duration persisted despite the reduction in myocardial stiffness.

### 3.5 Aged slices show pathway-level transcriptional responses post-collagenase treatment

Bulk RNA sequencing was performed to determine whether ECM remodeling and the associated reduction in tissue stiffness were accompanied by changes in the myocardial transcriptional profile. PCA demonstrated clear separation between young and aged LMS, consistent with a pronounced age-associated shift in the global transcriptional profile (Fig. 5a, S4d). Relative to young LMS, aged tissue exhibited increased expression of immune-associated genes, including *Ccl6* and *Arg1*, as well as the senescence-associated gene Cdkn2a. Conversely, expression of the mitochondrial gene *mt-Nd6* was reduced in aged LMS (Fig. S4b).

**Figure 5.**
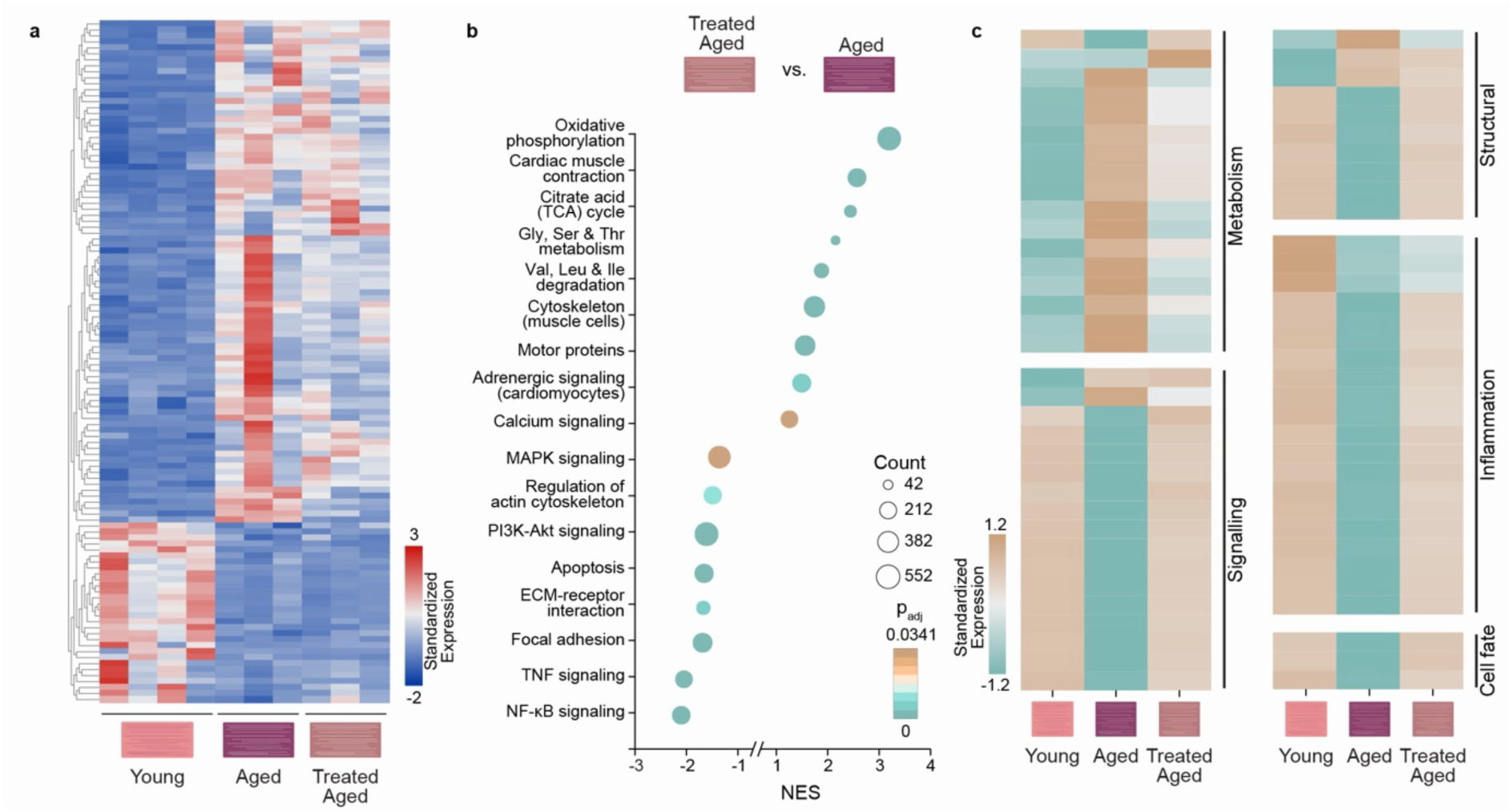
Transcriptomic changes associated with myocardial aging and collagenase treatment. **(a)** Heatmap of VST expression for all differentially expressed genes (DEGs) between aged and young LMS ((DESeq2; *p*_adj_ < 0.05, | log_2_ fold change | > 1 after ashr shrinkage), shown for all samples. Expression was row-standardized (*Z*-score), and rows were hierarchically clustered (Euclidean distance, complete linkage). **(b)** Gene set enrichment analysis (GSEA) of KEGG pathways in collagenase-treated aged versus untreated aged LMS. Genes were ranked by the DESeq2 Wald statistic across the entire detected transcriptome. Normalized enrichment scores (|NES|) > 1.5 indicate pathway enrichment.. Dot size represents the number of genes in each pathway (count), and color represents the Benjamini-Hochberg adjusted *p*-value (*p*_adj_ < 0.05). **(c)** Heatmaps of pathway-level normalized enrichment scores (NES) for KEGG pathways significantly enriched in the GSEA (*p*_adj_ < 0.05, |NES| > 1.5), grouped by overarching biological function. (*N* = 4 young, *N* = 3 aged, *N* = 3 collagenase-treated aged animals).

Collagenase-treated aged samples closely resembled untreated aged samples in the heatmap of age-associated differentially expressed genes (Fig. 5a). Consistent with this observation, treated aged and untreated aged LMS showed substantial overlap in the PCA space (Fig. S4d). No genes met the combined criteria for differential expression (Padj < 0.05 and |log2FC| > 1) in the comparison between collagenase-treated and untreated aged LMS (Fig. S4a). The comparisons of untreated and treated aged LMS with young LMS also shared 457 upregulated and 286 downregulated genes (Fig. S4f). Thus, although collagenase treatment produced marked changes in ECM composition and tissue mechanics, the global transcriptional profile remained characteristic of aged LMS.

To determine whether coordinated pathway-level changes were present, gene set enrichment analysis (GSEA) was performed using KEGG pathway gene sets. Aging was associated with enrichment of pathways related to signaling (including ECM-receptor interactions, focal adhesion, and NF-kB signaling) and cell fate (cellular senescence), whereas pathways associated with oxidative phosphorylation and cardiac muscle contraction were negatively enriched (Fig. S4e).

Comparison of collagenase-treated and untreated aged LMS revealed enrichment patterns in the opposite direction to aging for several of these pathways (Fig. 5c). Positively enriched pathways included oxidative phosphorylation, the tricarboxylic acid cycle, and amino acid metabolism and degradation. Cardiac muscle contraction and adrenergic signaling in cardiomyocytes were also positively enriched, together with calcium signaling. In contrast, ECM-receptor interactions, focal adhesions, PI3K-Akt signaling, MAPK signaling, apoptosis, TNF signaling, and NF-kB signaling were negatively enriched. Together, these findings indicate that ECM remodeling is accompanied by coordinated changes in selected transcriptional programs, while the overall expression profile remains characteristic of aged LMS.

## 4. Discussion

This study investigated whether age-associated remodeling of the cardiac ECM could be modified to alter myocardial mechanics and function. Using an *ex vivo* LMS platform, we found that controlled enzymatic remodeling of the aged ECM reduced collagen abundance and network complexity and shifted myocardial stiffness toward young tissue values. However, age-associated differences in contractility and the global transcriptional profile persisted over 48 hours. ECM remodeling was also accompanied by a higher proportion of regular beats and coordinated transcriptional changes in selected metabolic, cardiac, and inflammatory pathways. Together, these findings suggest that the aged myocardial ECM remains amenable to targeted structural remodeling, with distinct effects on tissue mechanics and biological responses.

Engineered *in vitro* ECM systems have shown that matrix age influences cardiomyocyte phenotype [35], while ECM ligands and stiffness distinctly regulate cardiac fibroblast activation [6]. This study extends the investigation of these relationships to native aged myocardial tissue, where resident cells remain embedded within their matrix. Collagenase has previously been used to investigate the contribution of collagen to ventricular geometry and mechanics in isolated hearts [42]. Here, direct remodeling of an established aged matrix was combined with assessment of structural, mechanical, contractile, and transcriptional responses during short-term culture. This approach allows the consequences of matrix perturbation to be examined within the multicellular tissue environment and provides a complementary model to systems that independently vary matrix composition and stiffness.

Aged myocardium exhibits remodeled ECM at both compositional and architectural levels. In aged LMS, increased collagen abundance, estimated fiber width, and network complexity were shown. Age-related increases in collagen content and fibril diameter have been reported in human myocardium [4]. Collagenase treatment reduced both collagen content and network complexity, while estimated fiber width did not differ significantly between treated and untreated aged LMS. No treatment-associated changes were detected in fibronectin metrics examined. Collagen network mechanics are influenced not only by the amount of collagen present, but also by network connectivity, fiber organization, and the manner in which load is distributed across the network [61–63]. The accompanying reduction in myocardial stiffness is consistent with altered load transmission, although the individual contributions of collagen loss and architectural remodeling could not be separated.

Another important finding of this work was that remodeling of the aged ECM reduced myocardial stiffness without a detectable change in ultimate tensile stress. Different mechanical properties can respond differently to enzymatic matrix remodeling. For example, collagen removal from the porcine heart valve leaflets altered stress relaxation without a significant change in biaxial extensibility [64]. In aged LMS, treatment reduced the linear modulus, while strain and energy at ultimate stress remained lower than in young LMS. Reduced stiffness therefore coexisted with persistent differences in deformation and energy absorption before maximum stress. This distinction is relevant to ECM engineering because a shift in stiffness toward young tissue values does not establish recovery of the broader mechanical phenotype.

Despite substantial evidence that ECM biophysical and biochemical cues affect cell behavior [6,33], remodeling of the aged ECM did not restore contraction duration to young tissue values. Normalized contraction amplitude and frequency did not different significantly among groups, while a higher proportion of regular beats was observed after treatment. A reduction in stiffness also modifies only part of the aged extracellular environment. For example, in DECIPHER scaffolds, young ECM ligands supported a quiescent cardiac fibroblast phenotype even at stiffnesses representative of aged myocardium [6], indicating that mechanical and biochemical matrix cues can have distinct effects on cell state.

The persistence of age-associated contractile features may reflect the timescale and nature of the intervention presented in this study. The two-day culture period may have been sufficient to remodel the established aged ECM but insufficient for downstream adaptation of the cellular phenotype. In addition, prolonged exposure to the aged ECM may lead to persistent cellular responses to prior mechanical cues (i.e., mechanical memory) [32,65]. Intrinsic alterations in calcium handling and mitochondrial function also accompany cardiac aging and may persist after extracellular cues are altered [66]. Whether these processes constrain the response of aged LMS to matrix remodeling remains to be determined.

Although treated aged LMS remained globally similar to untreated aged tissue, pathway-level analysis indicated that the aged myocardium exhibited a molecular response to treatment. No genes met the combined significance and fold-change threshold for differential expression between the two aged groups. Nevertheless, coordinated changes were observed in pathways associated with cardiac contraction and calcium handling, cellular metabolism, mechanotransduction, and inflammatory signaling. Positive enrichment of oxidative phosphorylation and cardiac muscle contraction, together with negative enrichment of ECM-receptor interactions, focal adhesions, and inflammatory signaling, opposed several changes observed with aging. These findings suggest that matrix remodeling can influence selected transcriptional programs while the broader aged expression profile persists. The enrichment results identify candidate cellular responses but do not establish recovery of the corresponding metabolic or contractile functions. Bulk RNA-seq also cannot distinguish changes within particular cell types from changes in their relative contributions to the tissue sample.

Several limitations should be considered when interpreting these findings. Static stretch during culture and electrical pacing during measurement do not reproduce continuous electromechanical conditioning, which can influence myocardial slice phenotype [44,47]. Further, the culture medium and oxygenation conditions differ from the native myocardial environment and may influence cellular metabolism and fibroblast activity. In addition, collagenase modifies matrix composition and architecture together with tissue mechanics, preventing attribution of the biological responses to stiffness alone. Possible contributions from altered ligand presentation or collagen degradation products remain unresolved.

Future studies should determine whether prolonged exposure to a remodeled ECM, dynamic electromechanical conditioning, or combining with interventions targeting cardiomyocyte metabolism, calcium handling, or cellular senescence can improve contractile function over longer timescales. Local measurements linking ECM architecture and mechanics with cell-specific responses would help identify which matrix changes are functionally consequential. The immediate value of this approach is as an experimental model for testing matrix-directed interventions in living aged tissue. Defining which features of the aged myocardium remain responsive to ECM remodeling will help determine how extracellular and cellular interventions can be combined to improve tissue function.

## 5. Conclusion

This study demonstrates that collagenase treatment remodels the aged cardiac ECM in *ex vivo* LMS, reducing collagen and network complexity, and shifting myocardial stiffness toward young tissue values. Despite these substantial extracellular changes, age-associated differences in contractile function and the global transcriptional profile persisted over 48 hours. Treatment was accompanied by pathway-level transcriptional changes involving metabolism, mechanotransduction, and inflammatory signaling. Together, these findings highlight the potential of targeted ECM remodeling to modify the aged cardiac microenvironment while showing that structural and mechanical changes do not necessarily produce a corresponding shift toward young tissue function. The LMS approach provides a means to evaluate how matrix-directed interventions influence mechanical and biological responses within native aged myocardium.

## CRediT authorship contribution statement

**Ranmadusha Hengst:** Conceptualization, Methodology, Investigation, Formal analysis, Data curation, Visualization, Writing (original draft). **Erik Spaans:** Methodology, Formal analysis. **Md. Faris H. Ramli:** Methodology, Formal analysis. **Avery Sun:** Methodology, Formal analysis. **Vicky Lai:** Methodology. **Kim Whye Leong:** Methodology. **Yinling Zhu:** Methodology. **Roger Foo:** Resources. **Jin Zhu:** Formal analysis. **Chii Jou Chan:** Resources. **Leo Hwa Liang:** Resources. **Jennifer L. Young:** Conceptualization, Supervision, Funding acquisition, Project administration, Writing (review & editing).

## Ethics declaration

This study was conducted in accordance with institutional guidelines for animal welfare and/or reporting. Tissues were obtained through the NUS Animal Tissue Sharing Program and used in accordance with approved protocols.

## AI declaration statement

During the preparation of this work, the authors used ChatGPT, Gemini, and Claude in order to provide coding assistance, feedback on content organization, and improving grammar and readability of the manuscript. After using these tools/services, the authors reviewed and edited the content as needed and take full responsibility for the content of the published article.

## Declaration of competing interests

The authors declare no completing interests.

## Acknowledgements

The authors thank the following people who contributed to this work: Jennifer Marlena (MBI, NUS) for technical assistance and scientific illustrations; Anwesha Guru and Raageshwari Arulselvam (MBI, NUS) for imaging assistance; Ong Huan Ting, Apoorva Shivankar, Boon Heng, and Shen Xingyu (MBI, NUS) for animal procedures and tissue collection; Rong Li and Chii Jou Chan (MBI, NUS) for providing mouse heart tissues via the NUS Tissue Sharing Program; Apoorva Shivankar and Chii Jou Chan for providing vibratome training and usage; Kelly Tan for assistance with holotomography imaging; Li Hangfeng and Pakorn Tony Kanchanawong for providing the pulse generator used for the electrical pacing system; Ong Hui Ting (MBI, NUS) for assistance with image analysis; Matthew Ackers-Johnson (CVRI, NUS), Jie Yan, Alexander Bershadksy, Alphée Michelot, and Young Lab members (MBI, NUS) for fruitful discussions and technical advice. We thank the MBI Wet Lab Core for facility and technical support and the Singapore Microscopy and Bioimaging Analysis (SiMBA, MBI) core for microscopy and data processing facilities

## Funding

This work was supported by the National Research Foundation, Singapore, Mid-sized grant NRF-MSG-2023-0001, Singapore Ministry of Education under the Research Centres of Excellence Programme through the Mechanobiology Institute and the Department of Biomedical Engineering at the National University of Singapore (NUS), as well as the Singapore Ministry of Education Academic Research Fund grants MOE-T2EP30125-0011 and MOET32021-0003.

## Data availability

The data supporting the findings of this study are available within the article and its Supplementary Information. RNA-seq data will be available on GEO. Raw images are available from the corresponding author upon reasonable request.

## Code availability

The codes used in the analyses will be publicly available on GitHub.

## Notes

### Competing Interest Statement

The authors have declared no competing interest.

