## Supplemental Data for "Enzymatic extracellular matrix remodeling in *ex vivo* slices of aged myocardium"

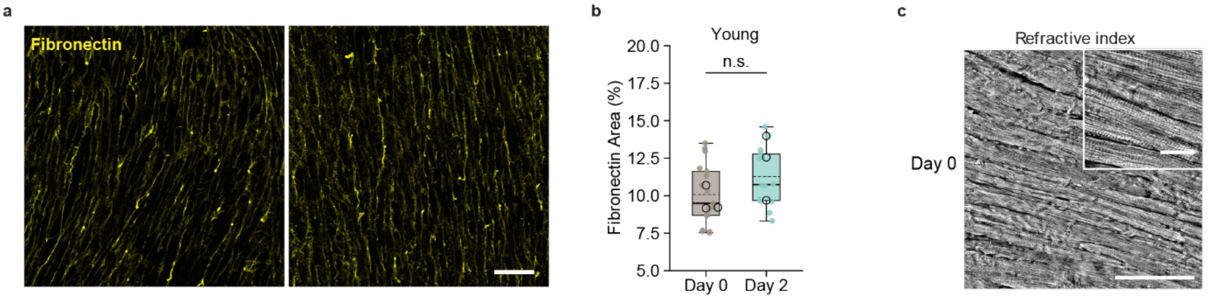


**Supplementary Figure 1. Fibronectin characterization and tissue morphology in young LMS (a)** Representative maximum intensity confocal projections showing fibronectin immunofluorescence in young LMS at baseline (Day 0, left) and after culture (Day 2, right). **(b)** Quantification of fibronectin-positive area via automated thresholding (%). Box plots display the median (solid horizontal line), mean (dashed horizontal line), and interquartile range (IQR; box bounds). Whiskers extend to the most extreme data points within 1.5 × IQR of the box edges. Filled points represent individual tissue positions ($n = 4 - 5$ fields of view per tissue); open circles indicate per-animal medians ($N = 3$ animals per group; LMM with Holm-Bonferroni correction, n.s. $p \geq 0.05$). **(c)** Representative holotomographic overview image of young LMS on Day 0 with high magnification inset. Scale bars = 100 $\mu$m (**a, c,** main) and 20 $\mu$m (**c,** inset).

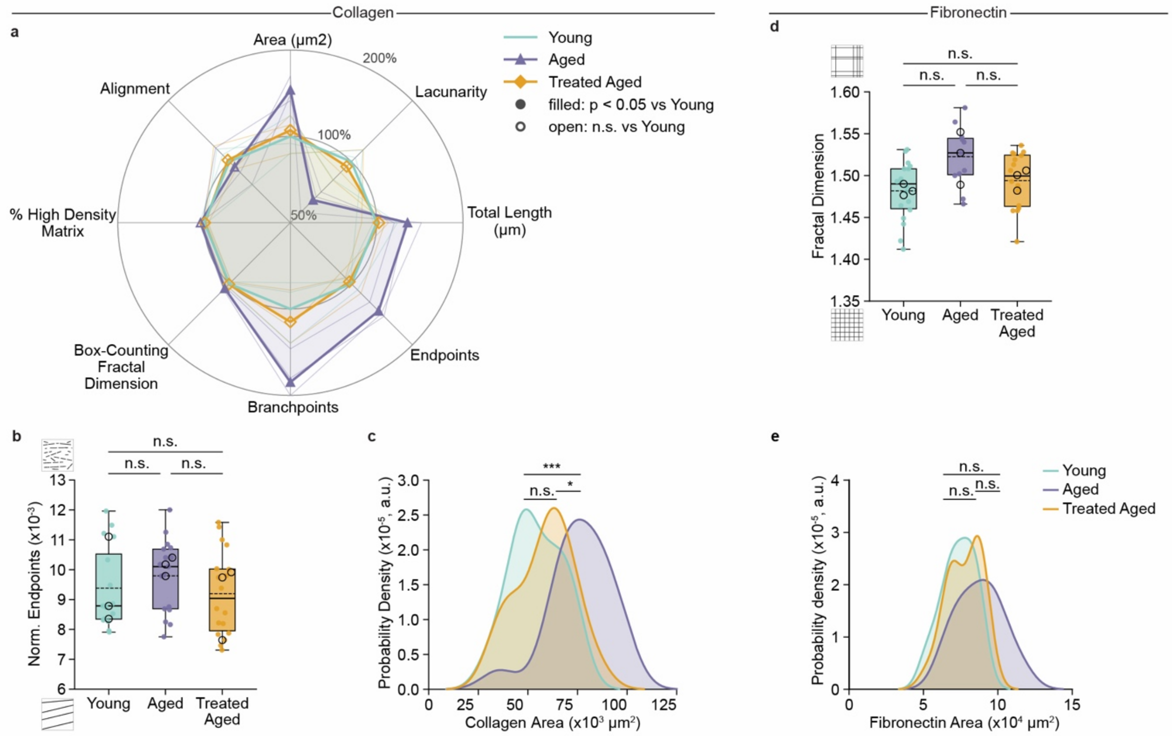


**Supplementary Figure 2. ECM architecture analysis in LMS. (a)** Radar plot comparing collagen architectural parameters relative to young controls (Young = 100%, grey circle; $\log_2$ radial scale). Filled markers denote $p < 0.05$ vs. Young; open markers indicate n.s. **(b, c)** Collagen matrix quantification showing (**b**) collagen matrix endpoints normalized to total fiber length and (**c**) probability density distributions of collagen area. **(d, e)** Fibronectin matrix quantification showing (**d**) box-counting fractal dimension and (**e**) probability density distributions of fibronectin area. Density distributions **(c, e)** are presented as normalized kernel density estimates (Probability density, a.u.). Box plots display the median (solid horizontal line), mean (dashed horizontal line), and interquartile range (IQR; box bounds). Whiskers extend to the most extreme data points within 1.5 × IQR of the box edges. Filled points represent individual tissue positions ($n = 5 - 7$ fields of view per tissue); open circles indicate per-animal medians ($N = 3$ animals per group). Pairwise group comparisons evaluated using LMM with animal as random intercept (Wald $z$), with $p$-values adjusted via Holm-Bonferroni correction ($* * * p < 0.001$, $* * p < 0.01$, $* p < 0.05$, n.s. $p \geq 0.05$).

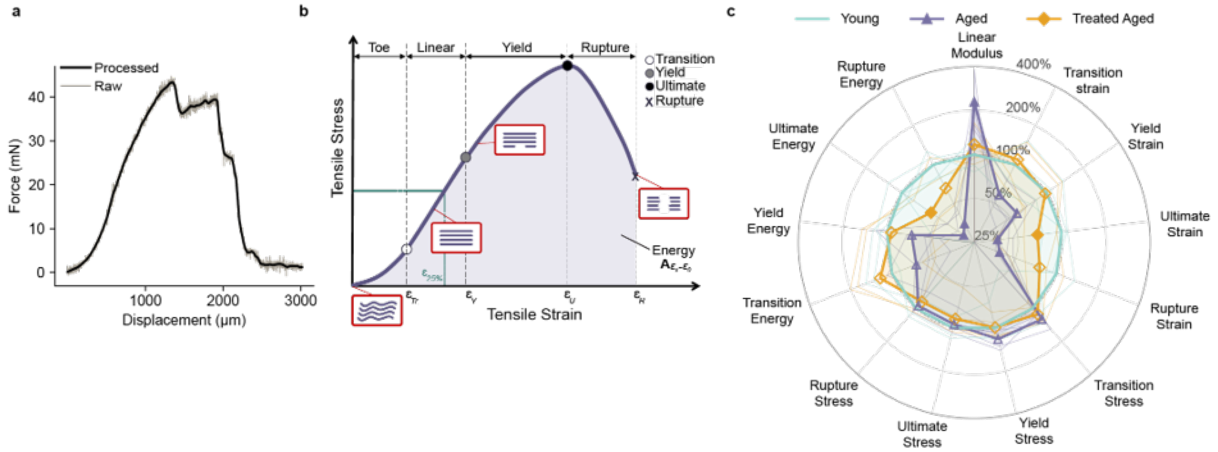


**Supplementary Figure 3. Biomechanical testing workflow, curve fitting, and regime definitions. (a)** Representative force–displacement curve demonstrating raw data and smoothed signal fitting. **(b)** Schematic illustration of mechanical testing regimes (toe, linear, yield, and rupture regions) alongside corresponding collagen fiber engagement and alignment states (uncrimping, linear recruitment/stretching, micro-failure, and ultimate rupture). **(c)** Radar plot comparing derived mechanical parameters relative to young controls (Young = 100%, grey circle; $\log_2$ radial scale). Filled markers denote $p < 0.05$ vs. Young; open markers indicate n.s. Statistical significance evaluated via LMM with Holm-Bonferroni post-hoc correction. ($N = 4$ young, $N = 5$ aged, $N = 4$ collagenase-treated aged animals; $n = 1 - 3$ tissue slices per animal). Abbreviations: $\varepsilon$, tensile strain; $\varepsilon_{Tr}$, transition strain; $\varepsilon_Y$, yield strain; $\varepsilon_U$, ultimate strain; $\varepsilon_R$, rupture strain; $\varepsilon_{25\%}$, 25% strain level; $A_{\varepsilon_x - \varepsilon_0}$, area under the stress–strain curve (energy strain density / toughness).

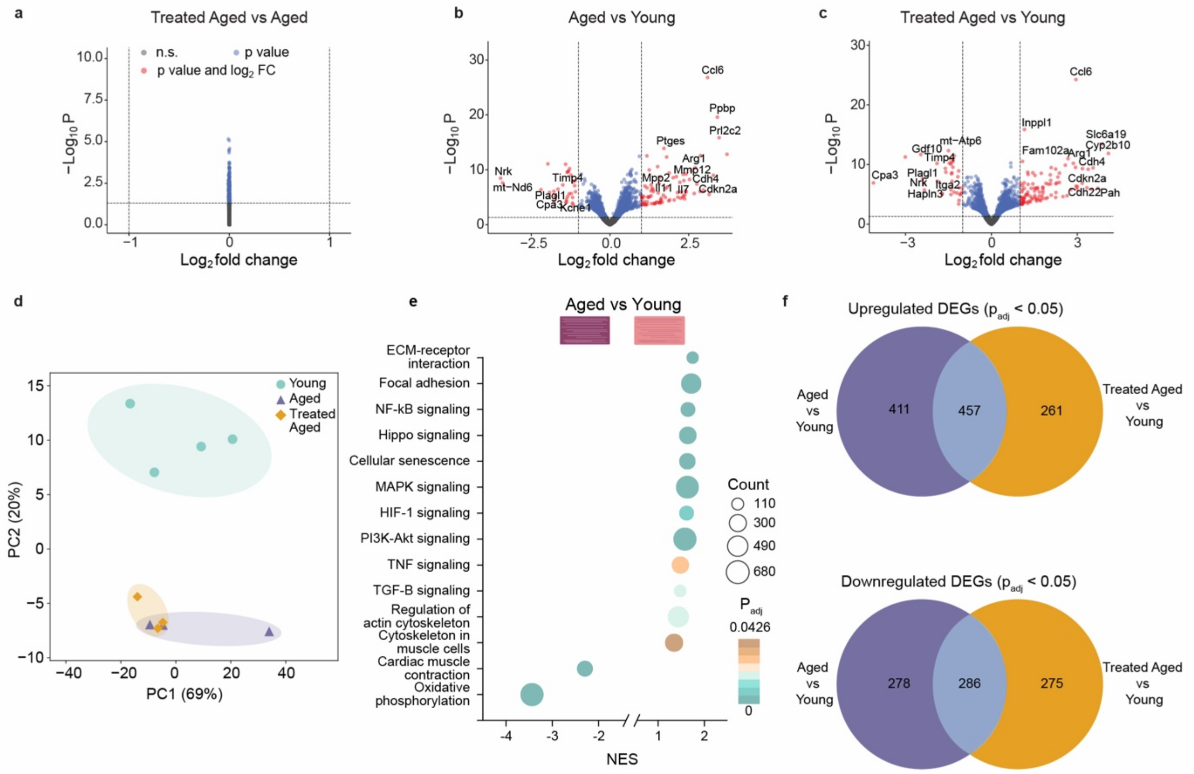


**Supplementary Figure 4. Differential gene expression and pathway enrichment analysis of LMS transcriptomes (a–c)** Volcano plots displaying differential gene expression for pairwise comparisons: (**a**) collagenase-treated aged vs. untreated aged, (**b**) aged vs. young, and (**c**) collagenase-treated aged vs. young LMS (DESeq2; $\log_2$ fold change shrunken using ashr estimator). Genes meeting significance threshold only ($p_{\text{adj}} < 0.05$) are shown in blue; genes meeting both significance and effect size thresholds ($p_{\text{adj}} < 0.05$ and $|\log_2$ fold change$| > 1$) in red; non-significant genes in grey. Horizontal and vertical dashed lines denote respective thresholds; select top DEGs are labeled. **(d)** Principal component analysis (PCA) of variance-stabilizing transformed (VST) bulk RNA-seq counts, showing separation of young from aged LMS along PC2 (20% of variance) and overlap between untreated aged and collagenase-treated aged LMS. **(e)** GSEA dot plot of enriched KEGG pathways in aged vs. young LMS performed across the transcriptome ranked by DESeq2 Wald statistic. Positive NES indicates enrichment in aged LMS; dot size represents gene count and color shade indicates adjusted $p$-value ($p_{\text{adj}} < 0.05$). **(f)** Venn diagrams showing shared and unique upregulated (top) and downregulated (bottom) DEGs ($p_{\text{adj}} < 0.05$) across experimental comparisons ($N = 4$ young, $N = 3$ aged, $N = 3$ treated aged animals).

**Supplementary Table 1: Antibodies and fluorescent probes used for LMS staining.**

| Target | Reagent | Supplier | Catalog No. | Dilution |
|---|---|---|---|---|
| Collagen | CNA35 | Addgene | 61603 | 1:60 |
| Fibronectin | Rabbit anti-fibronectin | Abcam | AB2413 | 1:100 |
| F-actin | Phalloidin CF568 | Biotium | 00044 | 1:250 |
| Nuclei | Hoechst 33342 | Invitrogen | H1399 | 1:2000 |
| Rabbit IgG | Goat anti-rabbit IgG (H+L), highly cross-adsorbed, CF®640R | Biotium | 20176 | 1:1000 |

**Supplementary table 2: Mechanical properties of young, aged, and collagenase-treated aged LMS derived from uniaxial tensile testing.** Values are presented as mean +/- standard deviation.

| | | Young | | Aged | | Aged Treated | |
|---|---|---|---|---|---|---|---|
| | | *Mean* | *Std.* | *Mean* | *Std.* | *Mean* | *Std.* |
| Linear (kPa) | Modulus | 78.6 | ± 20.8 | 192.0 | ± 77.8 | 102.7 | ± 30.8 |
| Strain | Transition | *0.3* | *± 0.1* | *0.1* | *± 0.0* | *0.2* | *± 0.1* |
| | Yield | 0.5 | *± 0.2* | *0.3* | *± 0.1* | *0.4* | *± 0.2* |
| | Ultimate | 1.3 | *± 0.3* | *0.5* | *± 0.1* | *0.8* | *± 0.3* |
| | Rupture | 1.6 | *± 0.5* | *0.6* | *± 0.2* | *1.1* | *± 0.4* |
| Stress (kPa) | Transition | *7.3* | *± 2.9* | *8.7* | *± 1.6* | *8.0* | *± 2.0* |
| | Yield | 23.7 | *± 5.8* | *28.0* | *± 4.9* | *24.0* | *± 5.1* |
| | Ultimate | 46.1 | *± 7.7* | *45.3* | *± 10.3* | *42.4* | *± 10.3* |
| | Rupture | 32.1 | *± 5.3* | *31.6* | *± 7.1* | *29.6* | *± 7.2* |
| Energy | Transition | *0.7* | *± 0.5* | *0.4* | *± 0.2* | *0.7* | *± 0.4* |
| | Yield | 4.2 | *± 1.5* | *2.5* | *± 1.2* | *3.8* | *± 2.2* |
| | Ultimate | 32.9 | *± 8.6* | *10.5* | *± 5.0* | *19.3* | *± 9.4* |
| | Rupture | 46.1 | *± 15.3* | *16.3* | *± 8.5* | *29.1* | *± 15.1* |
